# Non-neutralizing SARS-CoV-2 N-terminal domain antibodies protect mice against severe disease using Fc-mediated effector functions

**DOI:** 10.1101/2023.07.25.550460

**Authors:** Camille N. Pierre, Lily E. Adams, Kara Anasti, Derrick Goodman, Sherry Stanfield-Oakley, John M. Powers, Dapeng Li, Wes Rountree, Yunfei Wang, Robert J. Edwards, S. Munir Alam, Guido Ferrari, Georgia D. Tomaras, Barton F. Haynes, Ralph S. Baric, Kevin O. Saunders

## Abstract

Antibodies perform both neutralizing and non-neutralizing effector functions that protect against certain pathogen-induced diseases. A human antibody directed at the SARS-CoV-2 Spike N-terminal domain (NTD), DH1052, was recently shown to be non-neutralizing yet it protected mice and cynomolgus macaques from severe disease. The mechanisms of this non-neutralizing antibody-mediated protection are unknown. Here we show that Fc effector functions mediate non-neutralizing antibody (non-nAb) protection against SARS-CoV-2 MA10 viral challenge in mice. Though non-nAb infusion did not suppress infectious viral titers in the lung as potently as NTD neutralizing antibody (nAb) infusion, disease markers including gross lung discoloration were similar in nAb and non-nAb groups. Fc functional knockout substitutions abolished non-nAb protection and increased viral titers in the nAb group. Finally, Fc enhancement increased non-nAb protection relative to WT, supporting a positive association between Fc functionality and degree of protection in SARS-CoV-2 infection. This study demonstrates that non-nAbs can utilize Fc-mediated mechanisms to lower viral load and prevent lung damage due to coronavirus infection.

**AUTHOR SUMMARY:** COVID-19 has claimed over 6.8 million lives worldwide and caused economic and social disruption globally. Preventing more deaths from COVID-19 is a principal goal of antibody biologic and vaccine developers. To guide design of such countermeasures, an understanding of how the immune system prevents severe COVID-19 disease is needed. We demonstrate here that antibody functions other than neutralization can contribute to protection from severe disease. Specifically, the functions of antibodies that rely on its Fc portion were shown to confer antibody-mediated protection of mice challenged with a mouse adapted version of SARS-CoV-2. Mice given an antibody that could not neutralize SARS-CoV-2 still showed a decrease in the amount of infectious virus in the lungs and less lung damage than mice given an irrelevant antibody. The decrease in infectious virus in the lungs was even larger when the non-neutralizing antibody was engineered to mediate non-neutralizing effector functions such as antibody-dependent cellular cytotoxicity more potently. Thus, in the absence of neutralization activity, non-neutralizing binding antibodies can contribute to the overall defense against SARS-CoV-2 infection and COVID-19 disease progression.

## INTRODUCTION

COVID-19 has claimed over 6.8 million lives worldwide since it emerged in 2019 (1). In the United States, COVID-19 has become the third leading cause of death in adults (2) and the eighth leading cause of death in children and adolescents (3). The virus that causes COVID-19 disease, severe acute respiratory syndrome coronavirus-2 (SARS-CoV-2), mutates during its replication cycle, producing variants that escape immunodominant nAb responses elicited by vaccination or previous infection (4, 5). Thus, identifying other protective antibody functions to supplement the effects of neutralization is of particular importance in combating COVID-19 disease and future pandemics.

Non-nAb-mediated functions include antibody-dependent cellular cytotoxicity (ADCC), antibody-dependent cellular phagocytosis (ADCP), and complement-dependent cytotoxicity (CDC), which are mediated by the crystallizable fragment (Fc region) of an antibody (6). Antibody Fc-mediated effector functions are elicited in humans with COVID-19 (7, 8) and several correlative studies support that this immune response positively affects health outcomes. First, COVID-19 patients who recovered exhibited higher Spike-reactive antibody FcγR binding and antibody-dependent complement deposition than individuals who succumbed to disease (9). Second, correlative studies of human infection have suggested that individuals with more severe disease have a delay in antibody class-switching to IgG1 or IgG3, the emergence of serum RBD-specific antibody binding to FcγRIIa and III, and RBD antibody-dependent complement deposition or phagocytosis (9). Third, high vaccine efficacy after a single Spike mRNA immunization when nAb titers are low but binding antibody is high has supported the hypothesis that non-neutralizing antibodies (non-nAbs) may contribute to protection (10). Fourth, Spike mRNA immunization of mice lacking Fc gamma receptors (FcγRs) reduced vaccine protective efficacy against Omicron infection or heterologous betacoronaviruses (11, 12). Altogether, these studies support a role for antibody effector functions in protective SARS-CoV-2 immunity. However, two observations have obscured the role of antibody Fc effector functions in protecting against COVID-19 disease. First, antibody effector functions such as ADCP are higher in individuals who experience more severe disease (13). Second, the presence of Fc-mediated effector function activity is generally positively correlated with neutralization activity, making it difficult to delineate which antibody function affects disease outcome (14). Thus, the contribution of antibody Fc-mediated effector functions to protection from disease remains unclear.

N-terminal domain (NTD) neutralizing and non-neutralizing antibodies have been implicated as protective immune responses in active and passive immunization studies (15). In passive immunization studies in mice and nonhuman primates challenged with SARS-CoV-2, NTD non-nAb DH1052 reduced infectious virus titers, lowered lung hemorrhagic scores, lowered lung virus replication, and improved survival compared to control IgG-infused mice (15, 16). DH1052 interacted with mouse FcγRs and is hypothesized to bind the orthologous human FcγRs (15). This binding implicated Fc-mediated effector functions bridging the adaptive and innate immune systems to confer protection. Nonhuman primates infused with either NTD nAb DH1050.1 or NTD non-nAb DH1052 had suppressed viral subgenomic RNA to similar levels (15). Also, in nonhuman primates vaccinated with Spike NTD only and subsequently challenged with SARS-CoV-2, virus replication was suppressed to undetectable levels (16), despite only low serum neutralizing antibodies being detected (16). Serum antibodies elicited by NTD vaccination mediated NK cell degranulation (a marker of ADCC) (17), raising the possibility that non-nAbs contributed to the protection in the presence of the low nAb response in nonhuman primates.

The Fc of an antibody can be manipulated to alter affinity for FcγRs or complement protein C1q to determine the importance of Fc-mediated effector functions in protection from disease (18). The LALA-PG substitutions in the antibody Fc (L234A, L235A, P329G) are a well-established strategy for simultaneously eliminating binding to FcγRs and C1q in both mice and humans (19). In contrast, the DLE substitutions (S239D, A330L, I332E) increase antibody effector function, primarily through increased binding of Fc to the high affinity human FcγRIIIa (20). Thus, antibodies with DLE substitutions can be used to study changes in protective efficacy caused by enhancement of Fc functions. Previous studies with receptor binding domain (RBD) nAbs have used Fc knockouts to show Fc-mediated functions are important for protection by some RBD nAbs (21, 22, 23, 24). NTD antibodies have not been examined; thus, it is unknown whether increased or decreased Fc-mediated effector function activity results in differences in suppression of SARS-CoV-2 replication by non-neutralizing and neutralizing NTD antibodies.

Here, we hypothesized that Fc effector functions mediate protection by non-neutralizing antibodies, and substantially contribute to the protection afforded by neutralizing antibodies. We show that passive immunization with wildtype non-nAb DH1052 and nAb DH1050.1 was sufficient for protection against a mouse-adapted SARS-CoV-2 virus, though infectious virus titers in the lung were higher in non-nAb infusion versus nAb infusion. An Fc functional knockout version of non-nAb DH1052 possessing the LALA-PG substitutions did not protect against infection, indicating that Fc effector functions are necessary for non-nAb-mediated protection. Loss of Fc effector functions also increased viral titers in mice infused with nAb DH1050.1, though this increase did not result in a worse disease course. Finally, Fc enhancement using the DLE substitutions increased protection by non-nAb DH1052, confirming Fc-mediated protection from SARS-CoV-2 challenge in mice.

## RESULTS

### SARS-CoV-2 NTD antibodies with LALA-PG substitutions knockout mouse FcγR binding

To determine the role of Fc-mediated effector functions in protection against SARS-CoV-2 infection, we engineered wildtype and functional knockout versions of both non-neutralizing NTD antibody DH1052 and neutralizing NTD antibody DH1050.1 (**Figure 1A**). Starting with the sequence of human IgG1 (Fc allotype G1m17), we introduced the loss of function LALA-PG substitutions (L234A, L235A, P329G) (19). We produced the wildtype G1m17 and Fc knockout LALA-PG versions of DH1052 and DH1050.1 and tested their binding via ELISA, which confirmed introduction of the LALA-PG substitutions did not alter binding of the antibody to its target antigen (**Figure 1B**). We next used surface plasmon resonance (SPR) to determine binding of wildtype G1m17 and LALA-PG versions of DH1052 and DH1050.1 to mouse FcγRs I-IV (**Figures 1C-J**). Mouse FcγRs were tested because protective efficacy of these antibodies was planned to be assessed in mouse models of SARS-CoV-2 infection. To mouse FcγRs, LALA-PG substitutions eliminated detectable binding by both DH1052 and DH1050.1 (**Figures 1C-J**).

**Figure 1.**
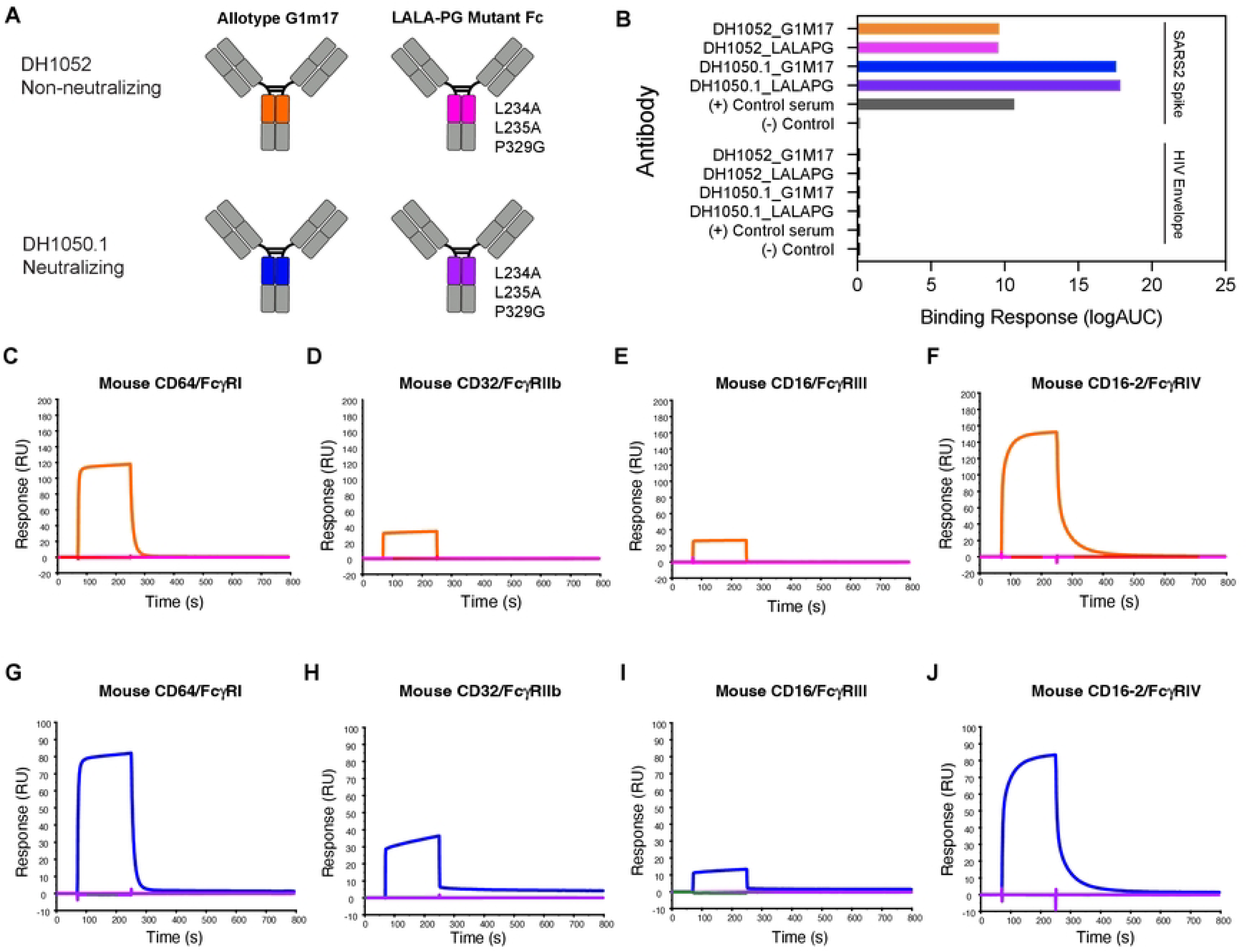
LALA-PG substitutions eliminate antibody binding to mouse FcγRI, II, III, and IV for both DH1052 and DH1050.1, without altering binding to SARS-CoV-2 Spike. (A) Antibody engineering schematic depicting wildtype (allotype G1m17) versus Fc-function knockout antibodies (LALA-PG substitutions: L234A, L235A, P329G). NTD-directed non-nAb DH1052 and nAb DH1050.1 are produced in both versions. Color scheme for each antibody is the same throughout A-J. (B) ELISA binding of G1m17 and LALA-PG antibodies to their cognate antigen SARS-CoV-2 Spike_D614G versus negative control antigen HIV-1 envelope. Binding response is measured as area under the log transformed curve (AUC). Serum from a nonhuman primate vaccinated with NTD was used as the positive control, and CH65 was used as the negative control antibody. (C-F) DH1052 G1m17 versus LALA-PG binding to immobilized mouse FcγRI, II, III, and IV measured via surface plasmon resonance (SPR). (G-J) DH1050.1 G1m17 versus LALA-PG binding to mouse immobilized FcγRI, II, III, and IV measured by SPR.

### NTD antibodies with enhanced FcγR binding show increased antibody-dependent cellular cytotoxicity (ADCC)

To further investigate the importance of Fc effector function, we also sought to up-modulate Fc effector functions. We engineered DH1052 and DH1050.1 to include DLE substitutions (S239D, A330L, I332E) (20), which are known to increase FcγR binding relative to wildtype Fc (**Figure 2A**). For both antibodies, DLE substitutions increased total binding to mouse FcγRs compared to wildtype G1m17 versions (**Figure 2B-I**), though FcγRI (**Figures 2B, 2F**) and FcγRIII (**Figures 2D, 2H**) peak binding was only modestly increased. Additionally, binding of DLE antibodies to FcγRI and FcγRIV showed slower dissociation compared to G1m17 (**Figures 2B, 2E, 2F, 2I**). Since DLE Fc modifications have been shown to increase ADCC mediated by FcγRIIIa in humans (20) and subtly increase ADCP (25), we assessed these two effector functions. ADCC activity in a natural killer (NK) cell degranulation assay using 293T cells expressing Spike D614G as a target showed CD107a surface expression, a marker of degranulation, was higher for DH1050.1_G1m17 compared to DH1052_G1m17 (**Figure 2J**). Both DH1052_LALA-PG and DH1050.1_LALA-PG did not trigger CD107a expression on NK cells (**Figure 2J**). Fc enhanced antibodies, DH1052_DLE and DH1050.1_DLE, generated substantial increases in NK cell CD107a expression compared to their G1m17 counterparts (**Figure 2J**). In total, the presence of DLE substitutions in the Fc were associated with increased ADCC activity.

**Figure 2.**
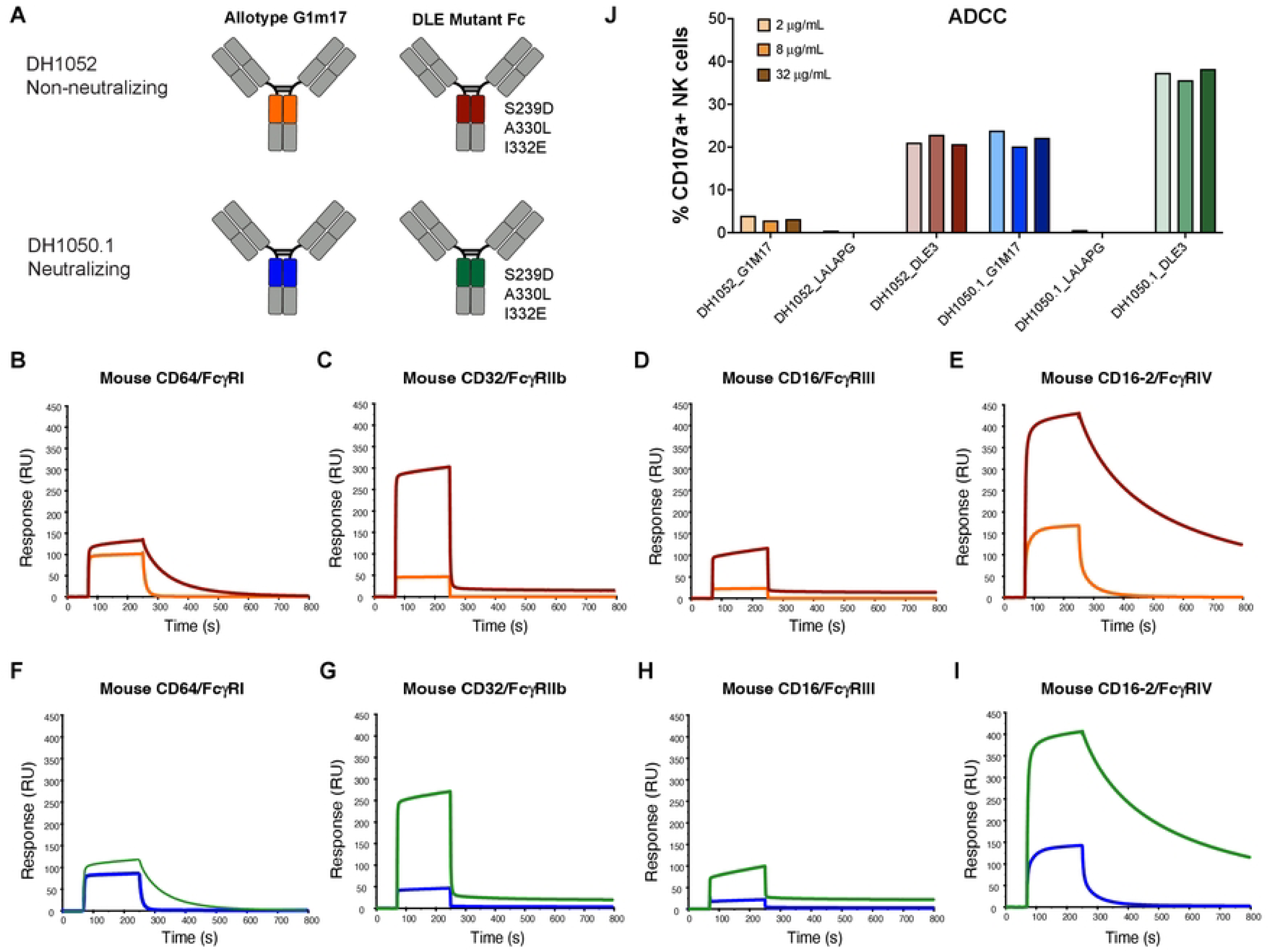
NTD antibodies with enhanced FcγR binding show increased antibody-dependent cellular cytotoxicity (ADCC). (A) Antibody engineering schematic depicting wildtype (allotype G1m17) versus Fc-function enhanced antibodies (DLE3 substitutions: S239D, A330L, I332E). (B-E) DH1052 G1m17 and DLE3 or (F-I) DH1050.1 G1m17 and DLE3 binding to immobilized mouse FcγRI, II, III, and IV measured by SPR. (J) Natural killer (NK) cell-mediated antibody-dependent cellular cytotoxicity of 293T cells expressing SARS-CoV-2 WA-1. Titers are shown as % NK cells expressing the degranulation marker CD107a. Each antibody was tested at 2, 8, and 32 μg/mL.

### LALA-PG substitutions eliminate or severely attenuate antibody-dependent cellular phagocytosis (ADCP)

We next compared the ability of G1m17, LALA-PG, and DLE versions of both antibodies to mediate antibody-dependent cellular phagocytosis (ADCP) of recombinant NTD. For use as a negative control antigen for non-neutralizing NTD antibodies, we designed a modified Spike NTD that eliminated binding of the non-neutralizing NTD antibodies. We solved the structure of DH1052 Fab in complex with the Spike trimer via negative stain electron microscopy (NSEM) and identified the loops at amino acids 70-76, 182-187, and 211-214 as candidate contact sites on the NTD (**Figure 3A and B**). We produced three NTDs that mutated each loop individually (NTD_ADEm1a-c) and one NTD that contained mutated versions of all three putative contact sites, NTD_ADEm3 (**Figure 3B**). Mutant NTD_ADEm1c and the combined mutant NTD_ADEm3 both eliminated binding of all members of a non-neutralizing NTD antibody panel (**Figure 3C**), indicating the critical loop for non-neutralizing NTD antibody binding includes the loop beginning at amino acid 211 (**Figure 3B**). Binding of neutralizing antibodies was not affected (**Figure 3C**), thus confirming the generation of a recombinant NTD that selectively knocked out non-nAb binding and consequently could be used for ADCP assays.

**Figure 3.**
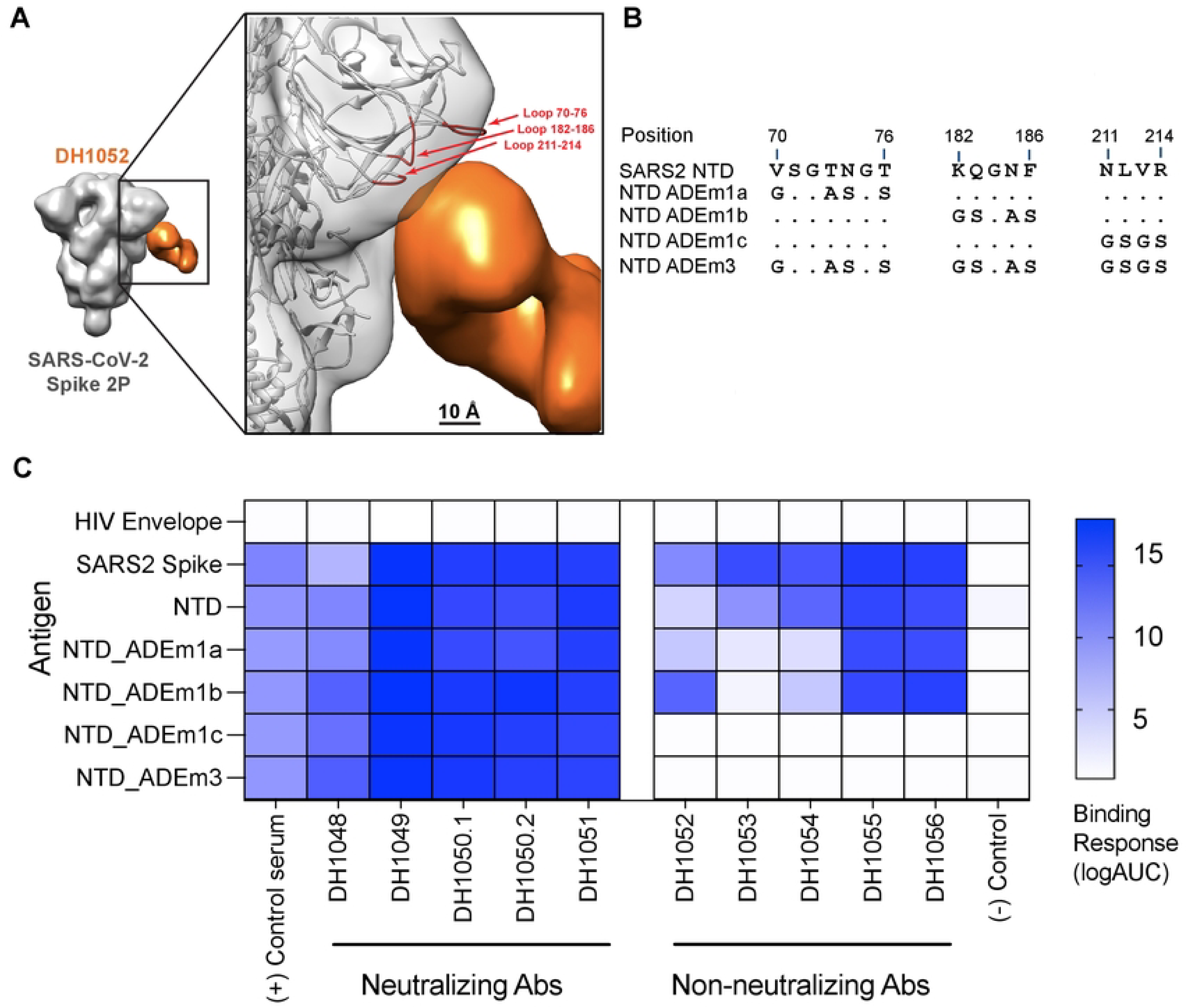
Modification of sites 211-214 in the Spike NTD eliminates binding by NTD non-nAbs. (A) Three-dimensional reconstruction by negative stain electron microscopy of DH1052 Fab (orange) in complex with SARS-CoV-2 Spike 2P (gray). In the enlargement, the density corresponding to the spike has been rigidly fit with a spike model (PDB 7QUS) shown in ribbon diagram. The Spike model is colored red where the NTD loops are most proximal to the Fab. NTD loops proximal to the putative DH1052 antigen combining site (red) were mutated as shown in B. (B) Sequence modifications for each mutant NTD tested. (C) ELISA binding results of a neutralizing NTD antibody panel (DH1048-DH1051, left) and a non-neutralizing NTD antibody panel (DH1052-DH1056, right) against mutant antigen candidates. NTD_ADEm3 combined the three mutations of NTD_ADEm1(a-c). Serum from a nonhuman primate vaccinated with NTD was used as the positive control, CH65 was used as the negative control antibody, and HIV Env was used as the negative control antigen.

Wildtype non-neutralizing DH1052_G1m17 incubated with ancestral Wuhan-Hu-1 SARS-CoV-2 NTD showed concentration-dependent ADCP activity, up to a point of saturation. At the highest concentration a prozone effect was observed where ADCP activity decreased (**Figure 4A**). No ADCP activity was observed for any version of DH1052 when the mutant NTD_ADEm3 was the antigen (**Figures 4A-C**). As expected, Fc knockout antibody DH1052_LALA-PG completely eliminated ADCP activity (**Figure 4B**). The ADCP score for DH1052_DLE was similar to DH1052_G1m17 consistent with published reports that DLE has subtle effects on ADCP (25)(**Figures 4C and 4D**). In concordance with binding reactivity, the neutralizing NTD antibody DH1050.1 mediated ADCP of both wildtype NTD and NTD_ADEm3 (**Figures 4E-H**). DH1050.1_G1m17 and DH1050.1_DLE showed similar ADCP activity (**Figure 4H**), whereas LALA-PG substitutions severely attenuated ADCP activity in DH1050.1 (**Figure 4F**). Thus, LALA-PG mutant antibodies exhibited severely reduced or ablated ADCP activity.

**Figure 4.**
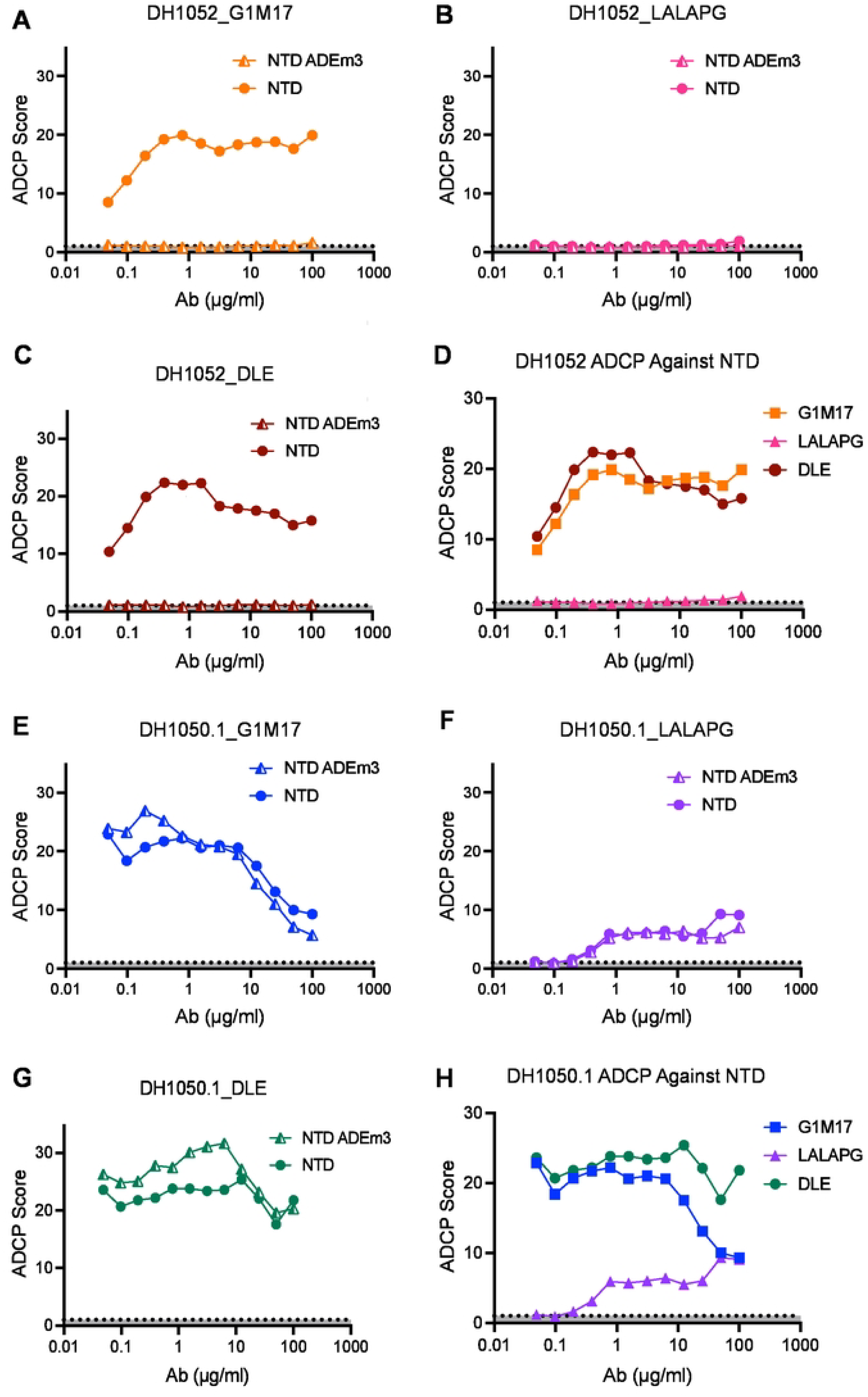
LALA-PG substitutions eliminate or severely attenuate antibody-dependent cellular phagocytosis (ADCP). (A-C) ADCP activity against NTD and mutant NTD_ADEm3 for G1m17, LALA-PG and DLE versions of DH1052 was determined in a THP-1 cell-based assay. Values shown are the mean of two technical replicates. ADCP score is a ratio of the fluorescence of the test result to the no antibody control (PBS). (D) A comparison of ADCP activity of WT NTD for all versions of DH1052 is shown. (E-G) ADCP of NTD and NTD_ADEm3 by G1m17, LALA-PG and DLE versions of DH1050.1 are shown. (H) A comparison of ADCP of Wuhan-Hu-1 NTD for all DH1050.1 versions is shown.

### DH1052_G1m17 and DH1050.1_G1m17 passive immunization protects *in vivo*

We hypothesized that Fc-mediated effector functions could protect against SARS-CoV-2 infection. BALB/c mice were given 300 μg of either test antibody (n = 10 per group) or isotype control antibody (n = 25) 12 hours prior to challenge with lethal SARS-CoV-2 MA10 virus (26) (**Figure 5A**). A group of five uninfected mice served as an additional negative control. Compared to isotype-treated mice, on day 4 post-infection, there was significantly less weight lost by mice that were administered either DH1052_G1m17 or DH1050.1_G1m17 compared to mice that received isotype control (**Figure 5B**; p<0.001, Wilcoxon test). Interestingly, mice administered DH1052_G1m17 showed a temporary mild weight loss on day 2 of infection, followed by a slight increase in body weight over the next 2 days. The initial decline in weight in mice given DH1052_G1m17 resulted in lower body weights in these mice compared to DH1050.1_G1m17-administered mice on day 4 (p<0.001). Infectious titers for mice administered either DH1052_G1m17 or DH1050.1_G1m17 were significantly lower than titers in mice given isotype control antibody (p<0.001) (**Figures 5C, 5F**). Gross lung discoloration scores for mice that received these antibodies were also significantly lower than in mice treated with the isotype control antibody (p<0.001), showing minimal congestion and hemorrhage after challenge (**Figures 5E, 5H**).

**Figure 5.**
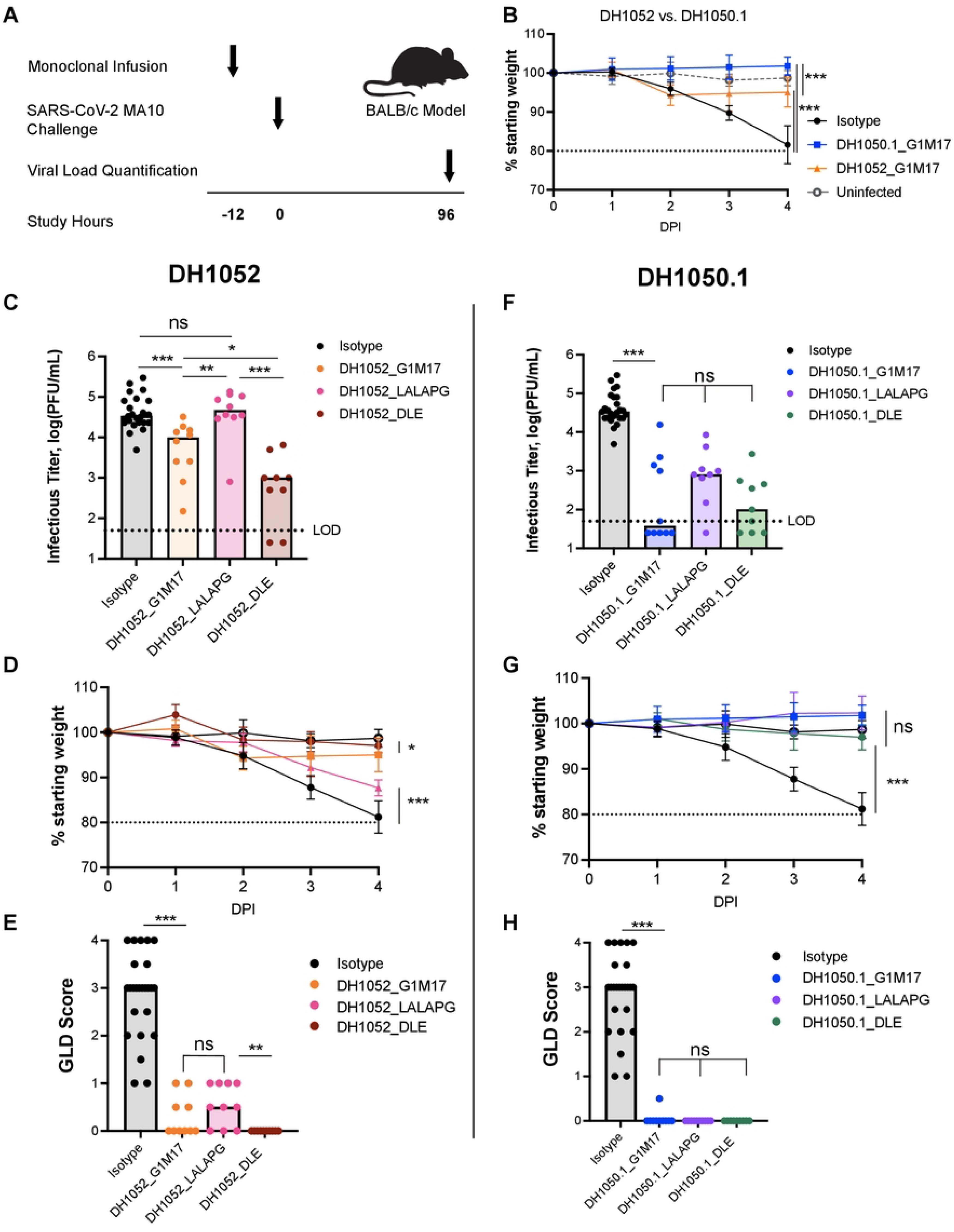
Fc knockout substitutions eliminated NTD non-nAb protection, and Fc enhancement increased NTD non-nAb protection. (A) Study design. BALB/c mice were passively infused at -12 hours and challenged with SARS-CoV-2 MA10 virus at 0 hours. Weights for each animal were collected each day of the experiment and lung tissue was harvested four days post-infection (DPI) to measure infectious viral titers and assign the gross lung discoloration (GLD) score. N = 10 mice per test group; n = 25 isotype control mice; n = 5 uninfected mice. (B) Percent weight loss in mice administered DH1052 G1m17, DH1050.1 G1m17, or Isotype control antibody. Uninfected mice were included as negative controls for weight loss. (C-E) Non-nAb DH1052 and (F-H) nAb DH1050.1 protection against infection and disease. Protection was assessed by (C,F) lung viral titers quantified as the log(PFU/mL), (D,G) weight loss each day post-infection (DPI) expressed as % original weight, and (E,H) median gross lung discoloration scores. All statistical comparisons were calculated using exact Wilcoxon rank sum tests using an alpha level of 0.05 (* p<0.05; ** p<0.01; *** p<0.001).

### Passive immunization with LALA-PG antibodies yielded higher infectious virus titers compared to wildtype antibody in mice

We next assessed whether LALA-PG versions of NTD non-nAbs or nAbs would diminish protection. Day 4 infectious virus titers in the lungs of mice treated with DH1052_LALA-PG matched titers of mice given isotype control antibody and were significantly increased compared to levels in mice infused with DH1052_G1m17 (p<0.001) (**Figure 5C**). Body weights steadily declined in the DH1052_LALA-PG group until the conclusion of the study on day 4 (**Figure 5D**). Unexpectedly, there was not a significant increase in macroscopic lung discoloration in the LALA-PG group compared to G1m17, despite the overall worse phenotype of the DH1052_LALA-PG group (Wilcoxon Test) (**Figure 5E**). Nonetheless, these results showed that loss of Fc-mediated effector functions led to loss of protection by non-nAb DH1052. The phenotype of neutralizing DH1050.1 was not altered as substantially by the LALA-PG Fc knockout substitutions, perhaps due to its neutralizing activity. Neither median infectious virus titers, body weights, or lung discoloration scores were significantly different between mice administered DH1050.1_G1m17 or DH1050.1_LALA-PG (**Figures 5F-5H**).

We sought to understand why LALA-PG mutant antibodies did not result in higher gross lung discoloration scores despite higher virus replication. An overabundance of proinflammatory cytokines in the lungs has been suggested to be one mechanism by which COVID-19 lung damage occurs (27). Thus, we compared the lung cytokine profile four days after SARS-CoV-2 MA10 challenge in mice that received LALA-PG or wildtype NTD antibody treatments. We quantified 26 cytokines in clarified lung homogenates and normalized the cytokine concentration to total protein concentration in the homogenate. When analyzed as fold change in normalized cytokine concentrations of each NTD antibody test group (n = 10 per group) compared to the isotype control group (n = 25; **Figure 6)**, we found differences in cytokine expression profiles between the wildtype and LALA-PG versions of non-neutralizing DH1052 (**Figure 6A**). Whereas proinflammatory cytokine IL-6 level was not significantly different between wildtype DH1052 and isotype control, the DH1052_LALA-PG group showed a significant decrease in IL-6 expression compared to isotype (p<0.001) (**Figure 6A**). Additionally, the DH1052_LALA-PG administration markedly increased antiviral cytokines such as TNFα, IL-12, IL-1β, and IFNγ compared isotype control (p<0.001) (**Figure 6A**). DH1050.1_G1m17-administration did not increase these cytokines to the same extent (**Figure 6B**). Cytokines and chemokines associated with T cell responses (IL-2, RANTES, MIP1α and MIP2α) were also increased in the LALA-PG group compared to DH1052_G1m17 (**Figure 6A**). Overall, the cytokine response had a Th1 bias with IFNγ, TNFa, IL-1 and IL-12 being elevated. Altogether, the Fc functional knockout DH1052_LALA-PG led to an increase in antiviral cytokines relative to isotype control treatment *in vivo*, with the increase being far larger than what G1m17 induced (**Figure 6A**). This response suggests in the absence of Fc-mediated activity, robust antiviral cytokine activity was upregulated, and less IL-6 was released. This cytokine profile was associated with minimal macroscopic lung discoloration despite high infectious titers (**Figure 5C and 5E**).

**Figure 6.**
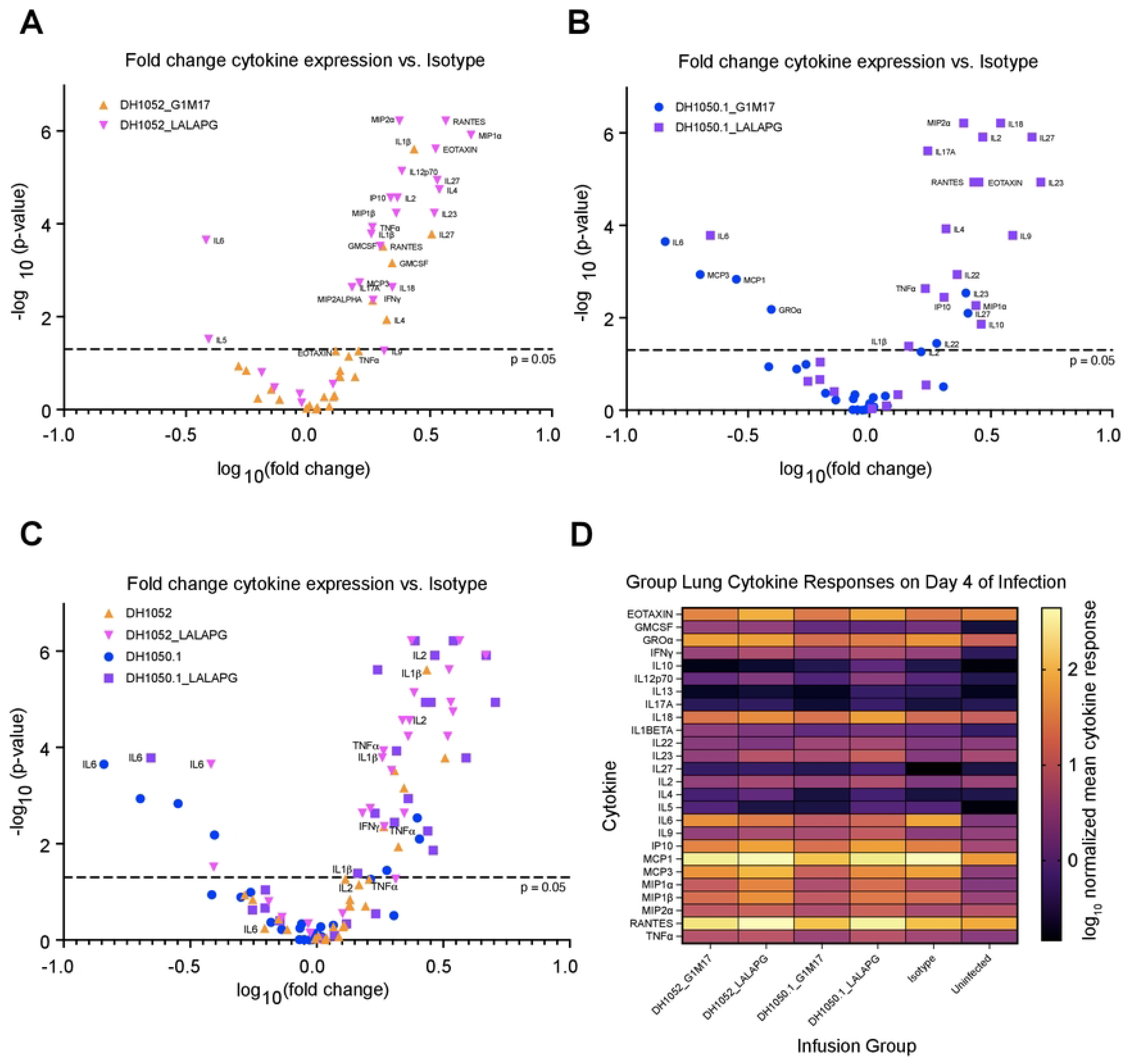
A distinct proinflammatory, antiviral cytokine response was elicited in mice after passive immunization with Fc knockout LALA-PG antibodies and challenge. Clarified lung homogenates obtained from mice day 4 post-infection were analyzed via Luminex multiplex assay for concentrations of 26 selected cytokines as labeled. The resulting concentrations (normalized for total homogenate protein) were compared to normalized concentrations in infected isotype control mice via Wilcoxon test. The fold change and p-value of each comparison between NTD antibody test group and isotype control are shown. (A) Comparisons between isotype control and the wildtype and Fc knockout DH1052 antibody versions. (B) Comparisons between isotype control and the wildtype and Fc knockout DH1050.1 antibody versions. (C) Overlay of all test groups as compared to isotype control. (D) Summary heatmap of mean normalized cytokine concentrations measured for each NTD antibody test group, infected isotype control, and the uninfected group of mice.

Comparison of wildtype and Fc-knockout versions of neutralizing DH1050.1 also showed marked differences between the two antibody treatments (**Figure 6B**). Either DH1050.1_G1m17 or DH1050.1_LALA-PG administration significantly lowered IL-6 lung concentration relative to isotype control (p<0.001) (**Figure 6B**). DH1050.1_G1m17 treatment also showed significantly lower MCP1, MCP3 and GROα (**Figure 6B**). As seen with DH1052_LALA-PG administration, DH1050.1_LALA-PG significantly increased a myriad of antiviral cytokines compared to infected isotype control mice (**Figure 6B and 6C**), but DH1050.1_G1m17 treatment did not. Moreover, the specific cytokines that increased expression were the same between DH1052_LALA-PG and DH1050.1_LALA-PG (**Figure 6C and 6D**). Therefore, eliminating Fc effector functions yielded the same alternate cytokine response for both neutralizing and non-neutralizing NTD antibodies.

### Fc-enhanced DH1052_DLE increased protection compared to wildtype DH1052

We hypothesized that if Fc effector functions mediated by DH1052 protected mice from SARS-CoV-2, then enhancing the ability of the antibody to mediate Fc effector functions would improve protection from SARS-CoV-2 infection. Our *in vitro* experiments showed that the DLE substitutions markedly increased ADCC and also modestly increased ADCP allowing us to test this hypothesis (**Figures 2J and 5**). We passively immunized mice and challenged them following the same protocol as with G1m17 antibodies (**Figure 5A**). DH1052_DLE showed significantly reduced viral titers compared to titers from DH1052_G1m17 mice (p = 0.016, Wilcoxon Test, n=10) (**Figure 5C**). The mice that received DH1052_DLE also had significantly higher final body weights compared to the G1m17 group (p = 0.01) (**Figure 5D**), and all mice in this group achieved lung discoloration scores of 0 (**Figure 5E**). Thus, enhancing the Fc function of a non-neutralizing NTD antibody improved protection from infection. Although DH1052 is a non-nAb and DH1050.1 is a nAb, there were no significant differences in infectious virus titers in the lungs (p=0.10), body weights (p=0.71), or lung discoloration scores (p=1.00) between groups of mice that received DLE versions of either of these two antibodies (**Figures 5C-H**).

Fc effector function enhancement did not result in major changes in protection by DH1050.1, owing to the fact that wildtype DH1050.1_G1m17 exhibited potent protection. Fc-enhanced DH1050.1_DLE showed no significant differences compared to DH1050.1_G1m17 lung infectious virus titers (p=0.30), body weights (p=0.85), or discoloration scores (p=1.00) (**Figures 5F-H**).

## DISCUSSION

With SARS-CoV-2 variants escaping from nAbs, the protection from severe disease afforded by non-nAbs is a key question. Even though current COVID vaccines no longer protect against transmission, they continue to protect against severe disease and death (28, 29, 30). Our study shows that wildtype non-nAbs can protect against manifestations of clinical disease in a mouse model. The mechanism of protection is Fc-mediated effector functions given that antibodies with ablated Fc effector functions conferred no benefit over negative control antibody. It should be noted that there were differences in the degree of protection by non-nAbs compared to nAbs. More specifically, mice that received non-nAb showed initial weight loss that subsided by day 2. This phenotype can be explained by the time it takes for initial infection of host cells to occur, viral antigen display on infected cells, and immune complex engagement by effector cells that clear infected cells. The initial weight loss demonstrates the different mechanisms neutralizing versus non-neutralizing antibodies use to control virus replication.

Loss of Fc effector functions diminished protection by the neutralizing NTD monoclonal antibody tested here. Our study focused on NTD-directed antibodies, but previous studies have examined receptor binding domain (RBD) antibodies. In agreement with our results for DH1050.1, multiple studies have shown that SARS-CoV-2 RBD-specific nAb protection can be dampened by loss of Fc-mediated functions(21, 22). Although, we should note that not all nAbs harness Fc effector functions to mediate protection (21, 22, 23). The difference in Fc effector function requirement for antibody-mediated protection has been attributed to differences in accessibility of the Fc region due to angles of approach (22, 31), but antibody neutralization potency, binding stoichiometry to Spike, and antibody epitope specificity may also explain the discordant results from different studies (6).

The antibody versions that lacked Fc engagement resulted in a large increase in cytokine secretion. As the lung faces invading pathogens continuously, the sources of lung cytokines has been intensely studied (32). This upregulation may reflect an increase in epithelial cell TNFα and IL-1β release in the presence of increased viral titers due to an absence of Fc-mediated innate cell activity (32). Epithelial cell cytokine responses may be combined with increased dendritic cell activation of T-cells given the observed significant increase in IL-2 after both DH1052_LALA-PG and DH1050.1_LALA-PG administration compared to isotype control (33). Overall, the cytokine response after Fc knockout non-nAb administration is most consistent with a Th1-biased response (IFNγ, TNFα, IL-12, IL-1). After infusion of the nAb lacking Fc effector function, both Th1 (TNFα) and Th2 responses (IL-4, IL-9) were observed (32, 34). Th1 responses are usually productive in resolving lung infections, and Th2 responses are associated with more severe lung dysfunction. We also found high concentrations of pro-inflammatory chemokines RANTES, MIP1α and MIP1β, which are secreted by CD8+ T cells (35). The overall immune response suggests that loss of Fc effector functions leads to a heterogeneous proinflammatory/antiviral cytokine response that is augmented relative to the natural responses to infection, resulting in less macroscopic lung damage than untreated infection.

IL-6 concentrations have been correlated with COVID-19 disease severity (36). Of the cytokines measured, only IL-6 was downregulated by both LALA-PG mutant antibodies compared to isotype control. We speculate that the infusion of LALA-PG versions of the antibodies could induce less IL-6 because immune complexes cannot engage FcγR on the surface of macrophages or monocytes (37). The low IL-6 seen in the LALA-PG groups would be expected to limit lung hemorrhage.

Initial concerns of antibody-dependent enhancement of SARS-CoV-2 infection by Fc-dependent attachment of antibody-virus immune complexes led to hesitancy in engaging Fc-mediated effector functions (38, 39, 40). These initial concerns led to therapeutic antibodies, such as Etesevimab being developed with mutated Fc regions that eliminate FcγR binding (41, 42). Here, we show Fc-mediated effector functions modulate viral load and disease severity in the absence of virus neutralization, indicating a clear benefit for Fc-mediated effector functions. Advances in Fc engineering and vaccine adjuvant design provide the technology to bolster Fc-mediated effector functions (25, 43, 44). The results of this study demonstrate Fc-mediated effector function can be beneficial for anti-coronavirus antibodies and for the next-generation of coronavirus vaccines.

We conclude here Fc-mediated functions are sufficient for protection against severe disease and lung damage in mice. The results presented here corroborate previous studies showing reduced betacoronavirus immunity after vaccination in FcγR knockout mice (11, 23). Antibodies that mediate ADCC arise during infection or vaccination and can be boosted by Spike mRNA vaccination in humans (10). We demonstrated that antibodies with increased ADCC capacity suppressed lung viremia better than wildtype antibodies, suggesting these antibodies may contribute to the protective efficacy of COVID-19 vaccines in the face of SARS-CoV-2 nAb escape (30). Non-nAbs can serve as a second line of defense where neutralization fails to protect due to immune evasion by the virus.

## METHODS

### Mouse Protection Studies and disease assessment

BALB/cAnNHsd were obtained from Envigo (strain 047) at 8-10 weeks of age on delivery where they were housed in groups under standard conditions. Twelve hours prior to infection, mice received 300 µg antibody or isotype control via intraperitoneal injection. Twelve hours after treatment, mice were infected with 10^4^ PFU SARS-CoV-2 MA10 (MA10) in 50 µl PBS intranasally under ketamine-xylasine anesthesia. Mice were weighed daily throughout the course of infection and were euthanized at 4-days post infection via isoflurane overdose for tissue collection and gross lung discoloration (GLD) scoring.

After euthanasia, lungs were scored for gross discoloration, indicating congestion and/or hemorrhage, based on a semi-quantitative scale of mild to severe discoloration covering 0 to 100% of the lung surface. The right inferior lung lobe was collected in 1 mL phosphate buffered saline with glass beads, homogenized, and debris was pelleted. Virus in the lungs was quantified from the inferior lobe suspension via plaque assay. Briefly, virus was serial diluted and inoculated onto confluent monolayers of Vero E6 cells, followed by agarose overlay. Plaques were visualized on day 3 post infection via staining with neutral red dye.

All cell lines and viruses were confirmed mycoplasma-negative, and viruses used were subjected to next-generation sequencing prior to use. Vero E6 cells were maintained in Dulbecco’s Modified Eagle’s Medium (DMEM) supplemented with 5% FBS and anti/anti-mouse adapted SARS-CoV-2 MA10 was developed based on the SARS-CoV-2 WA1 reference strain and propagated from a cDNA molecular clone (36, 37, 38).

All experiments were conducted after approval from the UNC Chapel Hill Institutional Biosafety Committee and Institutional Animal Care and Use Committee according to guidelines outlined by the Association for the Assessment and Accreditation of Laboratory Animal Care and the US Department of Agriculture. All Infections and downstream assays were performed at ABSL3 in accordance with Environmental Health and Safety. All work was performed with approved standard operating procedures and safety conditions for SARS-CoV-2. Our institutional ABSL3 facilities have been designed to conform to the safety requirements recommended by Biosafety in Microbiological and Biomedical Laboratories (BMBL), the US Department of Health and Human Services, the Public Health Service, the Centers for Disease Control and Prevention (CDC), and the National Institutes of Health (NIH). Laboratory safety plans have been submitted, and the facility has been approved for use by the UNC Department of Environmental Health and Safety (EHS) and the CDC.

### Recombinant antibody production

Recombinant antibodies were produced as described elsewhere (45). Heavy chain and light chain plasmids were obtained from GenScript. Expi293 cells (Life Technologies) were diluted to a final volume of 0.5L at a concentration of 2.5×10^6^ cells/mL in Expi293 media, and co-transfected with 400 μg each of heavy chain and light chain plasmids using Expifectamine. Five days after transfection, cell culture media was clarified by centrifugation and 0.8 μM filtration. Clarified culture media was incubated with Protein A resin overnight, washed with 25 mL of PBS containing 340 mM NaCl and eluted with 30 mL of glacial acetic acid. The pH of the eluted antibody solution was increased to neutral pH by adding 1M Tris pH8.0 and antibodies were buffer-exchanged in 25 mM Citrate, 125 mM NaCl, pH 6. Monomeric antibodies were purified by size exclusion chromatography on a Superdex 200 26/600 column (GE Healthcare) in 25 mM Citrate, 125 mM NaCl, pH 6, filtered, and stored at -80°C. All antibodies were confirmed Endotoxin negative using the Charles River Endoscan-V machine and program (software version 6.0.2).

### ELISA binding assay

384-well plates were coated with 2 μg/mL of antigens in 0.1 M sodium bicarbonate. Plates were stored at 4°C overnight and washed with PBS + 0.05% Tween-20 the following day. Blocking was performed with PBS + 4% (w/v) whey protein, 15% Normal Goat Serum, 0.5% Tween-20, and 0.05% sodium azide for 1 h at 25°C. After blocking plates were washed again with PBS + 0.05% Tween-20, and serial dilutions of antibodies or serum control were added. Antibodies were incubated at 25°C for 90 minutes and then plates were washed again. HRP-conjugated goat anti-human IgG secondary antibody (Southern Biotech) was used to detect binding in conjunction with TMB substrate (Sera Care Life Sciences).

### Surface Plasmon Resonance (SPR) binding assay

The SPR binding analysis of the COVID-19 monoclonal antibodies (mAbs) to recombinant mouse Fc-gamma receptors (FcγRs) was performed using a Biacore S200 or T200 instrument in HBS-EP+ 1X running buffer. Biotinylated mouse FcγRs (Sino Biological) were immobilized onto a Streptavidin sensor chip. Mouse CD64/FcγRI was immobilized to a level of approximately 100 RU; Mouse CD32/FcγRIIB and Mouse CD16/FcγRIII were immobilized to 350-400 RU; and Mouse CD16-2/FcγRIV was immobilized to approximately 150 RU. A blank Streptavidin flow cell (Fc1) was used as the negative control reference surface. The mAbs were tested at 100 μg/mL and were injected over the sensor chip surface for 180 s at 30 μL/min using the high-performance injection type followed by a 600 s dissociation. The mouse FcγR surfaces were then regenerated with one 12 s pulse of glycine pH 2.0 at 50 μL/min. Results were analyzed using the Biacore S200 or T200 Evaluation software (Cytiva). The blank streptavidin sensor surface along with buffer binding were used for double reference subtraction to account for non-specific protein binding and signal drift.

### Antibody-dependent cellular cytotoxicity NK cell degranulation assay

293T cells were cultured in DMEM (Gibco) supplemented with 10% heat-inactivated fetal bovine serum (FBS) and 100 μg/ml Penicillin and Streptomycin solution at 37°C and 5% CO_2_. Cell-surface expression of CD107a was used as a marker for NK cell degranulation, a prerequisite process for, and strong correlate of, ADCC (31), performed by adapting a previously described procedure (32). Briefly, target cells were 293T cells 2-days post transfection with a SARS-CoV-2 S protein G614 expression plasmid. Natural killer cells were purified by negative selection (Miltenyi Biotech) from peripheral blood mononuclear cells obtained by leukapheresis from a healthy, SARS-CoV-2-seronegative individual (Fc-gamma-receptor IIIA [FcγRIIIA]158 V/F heterozygous) and previously assessed for FcγRIIIA genotype and frequency of NK cells were used as a source of effector cells. NK cells were incubated with target cells at a 1:1 ratio in the presence of diluted monoclonal antibodies, Brefeldin A (GolgiPlug, 1 μl/ml, BD Biosciences), monensin (GolgiStop, 4 μl/6 ml, BD Biosciences), and anti-CD107a-FITC (BD Biosciences, clone H4A3) in 96-well flat bottom plates for 6 h at 37°C in a humidified 5% CO_2_ incubator. NK cells were then recovered and stained for viability prior to staining with CD56-PECy7 (BD Biosciences, clone NCAM16.2), CD16-PacBlue (BD Biosciences, clone 3G8), and CD69-BV785 (Biolegend, Clone FN50). Cells were resuspended in 115 μl PBS–1% paraformaldehyde. Flow cytometry data analysis was performed using FlowJo software (v10.8.0). Data is reported as the % of CD107a + live NK cells (gates included singlets, lymphocytes, aqua blue−, CD56+ and/or CD16+, CD107a+). All final data represent specific activity, determined by subtraction of non-specific activity observed in assays performed with mock-infected cells and in the absence of antibodies.

### Negative stain electron microscopy

Negative stain electron microscopy was performed as previously described (15). Fabs were prepared from IgG by digestion with Lys-C and mixed with recombinant spike protein at a 9:1 molar ratio. After incubation for 1 hour at 37 °C the mixture was diluted to 0.1 mg/ml with HEPES-buffered saline augmented with 5% glycerol and 7.5 mM glutaraldehyde, incubated for 5 minutes at room temperature, then 1 M Tris pH 7.4 stock was added to 75 mM final Tris concentration to quench excess glutaraldehyde. After quenching, a 5-µl drop of sample was applied to a glow-discharged carbon film on 300 mesh Cu grids, incubated for 10-15 seconds, blotted, stained with 2% uranyl formate and air dried. Grids were imaged with a Philips EM420 electron microscope operated at 120 KV at 82,000x nominal magnification and captured with a 2k x 2k CCD camera at a pixel size of 4.02 Å. Three-dimensional reconstructions were calculated with standard procedures using Relion 3.0 (46). Images were created using UCSF Chimera (47).

### Recombinant NTD Antigen production

The coronavirus proteins were produced and purified as previously described (15, 48, 49, 50). Plasmids encoding N-terminal domain proteins were obtained from GenScript. Plasmids were transiently co-transfected in FreeStyle 293-F cells using 293Fectin (ThermoFisher). All cells were tested monthly for mycoplasma. DNA was prepared using a Midiprep kit (Qiagen). On day 5 or 6 post transfection cell culture supernatants were clarified by centrifugation and filtration with a 0.8-μm filter. Stepwise purification included affinity chromatography using StrepTrap HP (Cytiva) ran in 1X Buffer W (IBA Lifesciences) and eluting in 1X Buffer E (IBA Lifesciences), and by size-exclusion chromatography using Superdex 200 columns (Cytiva) in 10 mM Tris pH8, 500 mM NaCl.

### Antibody dependent cellular phagocytosis (ADCP) assay

The ADCP assay was performed as previously described (51, 52, 53) with modifications. Briefly, quantification of ADCP was performed by covalently binding SARS-CoV-2 NTD and NTD_ADEm3 to NeutrAvidin fluorescent beads (ThermoFisher, Waltham, MA). Immune complexes were formed by incubation with serially diluted (2 fold) monoclonal antibodies DH1052 and DH1050.1 as wildtype, LALA-PG or DLE mutated versions (described above). Monoclonal antibody CH65 IgG1 served as a negative control (54). Immune complexes were incubated with THP-1 cells (ATCC, Manassas, VA), and cellular fluorescence was measured using a BD LSR Fortessa (BD Biosciences, San Jose, CA).

### Luminex Cytokine assay

Lung homogenate protein concentrations were determined using a Bio-Rad DC protein assay performed according to the manufacturer’s protocol and read on a Synergy H1 plate reader (Agilent). Concentrations were calculated by extrapolation from a BSA standard curve using Gen5 v.3.00 (Agilent). Cytokines in undiluted homogenate were quantified using a 26-plex Luminex bead array assay (ThermoFisher EPX260-26088-901) performed according to the manufacturer’s protocol and read on an Intelliflex DR-SE (Luminex Corp.). Cytokine concentrations were calculated by extrapolation from standard curves using Bio-Plex Manager v.6.2 (Bio-Rad). Concentrations of cytokines were divided by the corresponding total homogenate protein concentration and log_10_ transformed.

### Statistical Analyses

Exact Wilcoxon rank sum tests were performed using an alpha level of 0.05 to compare differences between groups using SAS (SAS Institute, Cary, NC). There were no corrections made for multiple comparisons. Mean values are plotted with standard deviation.

## AUTHOR CONTRIBUTIONS

C.N.P. and K.O.S. designed research strategy. C.N.P, L.E.A., K.A., J.M.P., D.G., S.S.O., D.L., and R.J.E. performed research experiments. C.N.P., L.E.A., S.S.O., W.R., Y.W., R.J.E analyzed data. G.F., G.D.T., R.S.B., B.F.H., and K.O.S. oversaw research and provided expertise during development. C.N.P. and K.O.S. wrote the manuscript with input from authors.

## ACKNOWLEDGEMENTS

We would like to thank Nolan Jamieson for assistance and quality control measures in reagent production. We would like to thank Robert Parks and Whitney Edwards Beck for organizing collaboration on passive immunization studies between Duke University and UNC Chapel Hill. We would like to thank Ande West for help with preparing and conducting mouse passive immunization studies. Biomarker profiling was performed under the management of Barbara Theriot and direction of Dr. Andrew N. Macintyre in the Immunology Unit of the Duke Regional Biocontainment Laboratory (RBL), which received partial support for construction from the National Institutes of Health, National Institute of Allergy and Infectious Diseases (UC6-AI058607). The work was also supported by a grant from the National Institutes of Health, National Institute of Allergy and Infectious diseases (P01AI158571).

